# Ocular Disease Prevalence in California Sea Lions In Human Care: Freshwater Versus Saltwater Housing

**DOI:** 10.1101/2024.12.20.629697

**Authors:** Ingrid Brehm, Silas Herzner, Katrin Baumgartner, Jörg Beckmann, Ralph Simon, Lorenzo von Fersen

**Affiliations:** Animal Physiology, Friedrich-Alexander Universität Erlangen-Nürnberg, Erlangen, Germany; Nuremberg Zoo, Nuremberg, Germany; Behavioral Ecology and Conservation Lab, Nuremberg Zoo, Nuremberg, Germany; Machine Learning and Data Analytics Lab, Friedrich-Alexander Universität Erlangen-Nürnberg, Erlangen, Germany

**Keywords:** California Sea Lion, Ocular health, Freshwater, Saltwater, Seasonal patterns

## Abstract

California sea lions (*Zalophus californianus*), are susceptible to various ocular diseases, with some literature suggesting a heightened risk for those kept in freshwater pools as opposed to saltwater. Given the potential implications of housing conditions on animal health, we wanted to analyze the ocular health of two distinct groups of sea lions kept at Nuremberg Zoo, one housed in freshwater and the other in saltwater. Data extracted from the animals’ medical records over a ten-year period were used to compare the incidence of eye conditions and other medical conditions observed. The results revealed no significant difference in the overall prevalence of eye diseases between the two environments. However, a distinct seasonal pattern was noted: sea lions kept in freshwater exhibited a peak in eye disorders during the summer months, while those in saltwater displayed a more uniform distribution of the occurrence of ocular diseases throughout the year. These findings suggest that sun exposure and water quality are potentially more influential factors in the development of ocular diseases in pinnipeds than salinity or UV radiation. Further studies are necessary to elucidate the underlying mechanisms and optimize care practices for these marine mammals.

**Simple Summary:** This study investigated the ocular health of California sea lions (*Zalophus californianus*) at Nuremberg Zoo, comparing those housed in freshwater pools to those in saltwater. An analysis of medical records over ten years showed no significant difference in the overall prevalence of eye diseases between the two groups. However, sea lions in freshwater experienced a peak of eye disorders in the summer, while those in saltwater had a more consistent occurrence throughout the year. These results imply that factors such as sun exposure and water quality may play a more crucial in ocular disease development than salinity or UV radiation. The study highlights the need for further research to understand these dynamics and improve care for marine mammals.

## 1. Introduction

Pinnipeds are commonly found in zoos and aquariums around the world. For example, 326 California sea lions (*Zalophus californianus*) are currently living at 64 institutions accredited by the EAZA (European Association of Zoos and Aquaria). Ensuring individual health is one of the most important factors in guaranteeing optimal welfare for animals in managed collections. Eye problems are common in these populations and often necessitate complex medical treatments and surgical interventions, which can be challenging. A prevalent opinion is that the absence of seawater in certain managed populations contributes to the development of eye diseases. However, it is important to consider that pinnipeds in the wild thrive in aquatic environments with varying salinity levels. While most pinniped species are primarily marine, several have exhibited remarkable resilience in freshwater habitats, either seasonally or even constantly. Notable examples include the Baikal seal (*Pusa sibirica*), which is uniquely adapted to live exclusively in Lake Baikal, Siberia, making it one of the few true freshwater seal species (Parsons, 2013). Furthermore, the ringed seal (*P. hispida*) has two notable subspecies that inhabit freshwater systems: the Ladoga seal (*P. hispida ladogensis*), found in Lake Ladoga in Russia, and the Saimaa ringed seal (*P. h. saimensis*), which resides in Lake Saimaa in Finland. Both subspecies are critically endangered. Additionally, the harbor seal (*Phoca vitulina*) has been observed in estuarine and riverine systems along the coasts of North America and Europe, often entering freshwater rivers to hunt for fish or haul-out on riverbanks (Smith, 2006). A population of harbor seals is resident to Iliamna Lake in Alaska (Hauser et al., 2008), a subspecies of the harbor seal in Canada, the Ungava seal (*P. v. mellonae*), lives exclusively in freshwater (Enns et al., 2020). The California sea lion (*Zalophus californianus*) and Steller sea lion (*Eumetopias jubatus*) have also been documented in freshwater environments, albeit temporarily, as they migrate between breeding sites and feeding areas, exploiting available resources (Harrison, 2018). Considering this adaptability to various aquatic environments, it can be assumed that also for sea lions in human care seawater is not strictly essential. The occurrence of eye problems in zoos and aquariums should therefore not only be attributed to water salinity, especially considering that pinnipeds have evolved to live in a variety of aquatic ecosystems. Understanding the multifaceted factors contributing to ocular health in these animals is crucial for improving their welfare in managed settings.

**Figure 1.**
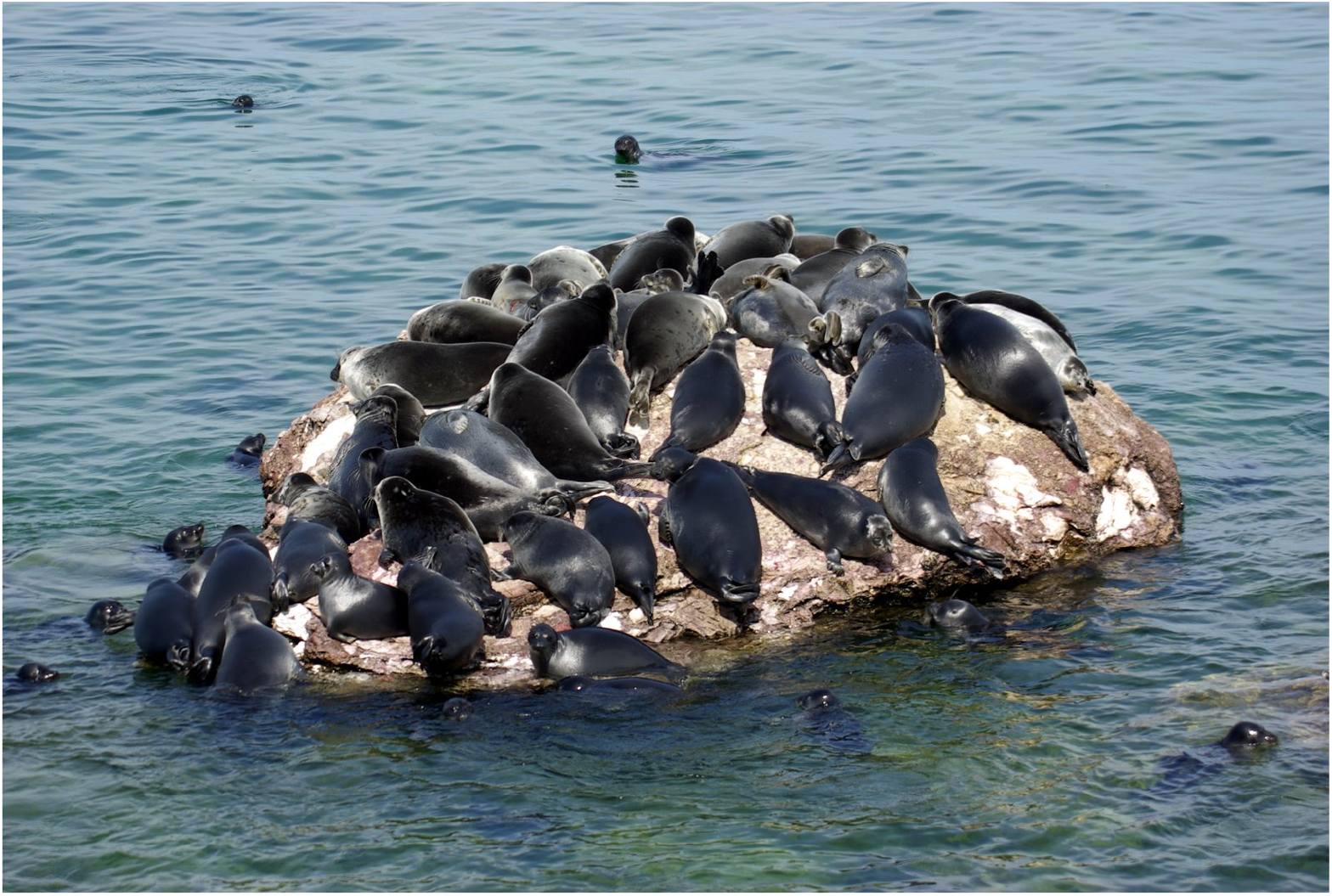
Group of Baikal Seals (*Pusa sibirica*) resting on a rock in Lake Baikal. This species lives exclusively in fresh water (credits: Sergey Gabdurakhmanov).

To assess the causes of eye problems in pinnipeds, it is essential to define the environmental challenges faced by these animals in human care compared to their wild counterparts (Gage 2011). Eye disease is one of the most common health problems in pinniped collections, and there is increasing evidence that solar radiation plays a significant role. Pinniped eyes are adapted to low-light environments, particularly for foraging at depth or in murky waters with minimal light penetration. In zoos, however, pinnipeds are often fed at the surface, exposing their eyes to much higher levels of solar radiation than they would typically experience in the wild.

In zoos, brightly painted pool walls are a significant factor in increased UV exposure, as they absorb UV radiation less effectively than darker surfaces. In addition, sea lions in human care frequently need to look upwards during training, feeding, and presentations, which can elevate their exposure to UV radiation. Certain enclosure designs necessitate that sea lions look up at keepers, contributing to this increased exposure. Additionally, heightened social interactions within the group can lead to more frequent eyelid opening, further enhancing their UV exposure (Gage 2011). Collectively, these factors increase the duration that these animals spend with their eyes open, thereby boosting their UV exposure levels (Nakamura et al., 2021). Colitz et al. (2010b) showed that ocular lesions in otariids were more common in months with higher sunlight intensity and longer daylight hours. In addition, facilities located closer to the equator report higher incidences of eye disease (Colitz et al., 2019). Even sunny winter months, when snow increases glare, can exacerbate the effect. Considering these findings, the German Federal Ministry of Food and Agriculture (BMEL) recommends using darker pool colors and providing shaded feeding areas to minimize sunlight exposure (BMEL 2020).

Keeping sea lions in freshwater pools has been identified by several specialists as a potential contributor to eye problems in pinnipeds. Some argue that housing sea lions in freshwater may lead to an increased risk of eye diseases, as these animals are naturally adapted to marine environments. Dunn et al. (1996) found that lense cataracts in harbor seals, were three times more common in freshwater than in saltwater. However, this has not been conclusively demonstrated for *Zalophus californianus*. Stach and Eule (2021) also found a correlation between freshwater housing and the development of corneal oedema in *Zalophus*. In their study of 209 pinnipeds in 25 facilities in Germany, Austria, and Switzerland, they reported that eye disease was significantly more common in animals housed exclusively in freshwater than in those with access to saltwater pools. The German Federal Ministry of Food and Agriculture (BMEL) recommends that pinnipeds should be housed in seawater or water with similar salinity (BMEL 2020). Colitz et al. (2019) further found that the occurrence of keratopathy (in %) was higher in housings with salinity levels below 29 g/L (64%) compared to salinity levels of over 29 g/l (52.7%) and concluded that a higher salinity provides some protection against this condition, and that keratopathy is more difficult to manage in animals housed in freshwater.

However, these findings have been challenged by research that investigated other pinniped species, which has shown that grey seals (*Halichoerus grypus*) can thrive in freshwater environments. A study in North Rona, Scotland, using Ecological Niche Factor Analysis, found that lactating female grey seals showed a clear preference for lower salinity pools. This behavior was most pronounced early in the season when thermal stress is highest, suggesting that grey seals may use freshwater pools not only for cooling but also for drinking. This evidence challenges the assumption that seawater is essential for the well-being of managed pinnipeds, suggests that freshwater may be a viable environment for some pinniped species, and questions whether salinity alone is responsible for the observed eye conditions (Stewart et al., 2014).

In support of this, Gage (2011) raised doubts about the effect of freshwater housing on eye disease, citing personal observations of sea lions in a northern freshwater facility with a black pool and a constant flow from an underground aquifer. None of the animals showed corneal lesions, in contrast to sea lions housed in a southern facility with natural saltwater but light blue, highly reflective pools, where many developed corneal damages. Gage (2011) argued that certain freshwater environments do not cause eye disease and that other factors, such as pool color and light reflection, may be more important.

A third factor mentioned in relation to eye problems in pinnipeds in human care is their exposure to oxidizing agents. The concentration of these agents in water systems can have a significant impact on the eye health of California sea lions. Halogens, primarily chlorine, or ozone, are commonly used to treat pool water. These halogens can react with organic matter, particularly nitrogen compounds, to form halogenated hydrocarbons or halogen amides, which are known to be irritating to the eyes. Halogen amides, such as mono- or dichloramines, can cause discomfort, while chlorinated hydrocarbons can cause liver damage (see Gulland et al. 2018). In addition, chlorinated hydrocarbons can interfere with the cytochrome P450 enzyme system in the ciliary body of the eye, potentially causing damage there as well (see Latson 2011).

Colitz et al. (2019) suggested that “chemicals used to disinfect enclosures and disinfection by-products may affect the preocular tear film and corneal epithelium, predisposing corneas to ulceration and secondary infection”. Further research is needed in this area. De Haan (2011) showed a correlation between the concentration of total, free and bound chlorine in swimming pool water and the severity of corneal lesions, with bound chlorine showing the most significant effect. The EAZA and EAAM (European Association for Aquatic Mammals) guidelines for the care of eared and fur seals recommend careful monitoring of water quality when using chlorine for disinfection and advise that the total chlorine concentration should not exceed 1.0 ppm (see Gulland et al. 2018).

When using ozone for water treatment, it is important to ensure that ozone is removed from the water as much as possible after its reaction in sealed chambers. The optimum ozone reduction potential (ORP) for treated water should exceed 700 mV, while water in contact with animals should maintain an ORP of less than 400 mV to prevent eye damage (Gulland et al. 2018). Higher ORPs in pool water can lead to conditions such as blepharospasm or epiphora (see Gage 2011). Interestingly, less prevalence of eye diseases is reported in pinnipeds housed in water without chemical additives (Nakamura et al. 2021). In addition, Colitz (2022) showed that the combination of ozone and chlorine can significantly promote the development of eye disease.

The fourth factor influencing eye problems in pinnipeds in human care is age. Like humans and dogs, age has a significant effect on the development of eye disease in Zalophus californianus. Colitz et al. (2010b) found nuclear sclerosis in animals aged 15 to 20 years and cataracts in all individuals over 26 years of age. They attributed these changes to a decline in antioxidant capacity with age (see Colitz et al. 2010b). Nakamura et al. (2021) found a correlation between age and lens disorders, noting that older animals had a higher incidence of such problems, while younger animals had more corneal lesions, probably due to increased activity leading to eye injuries. Colitz et al. (2019) confirmed that animals over 20 years of age had significantly higher rates of keratopathy.

The role of sex in predisposing to eye disease has also been discussed. Stach and Eule (2021) found that male pinnipeds more often suffer from eye diseases than females, a finding supported by Miller et al. (2013). However, Nakamura et al. (2021) found no correlation between the incidence of eye disease and sex. Aggressive behavior, particularly in males, may contribute to more eye injuries, as noted by Colitz et al. (2010b) and Nakamura et al. (2021).

Notable differences were observed in the distribution of eye diseases between wild individuals and those in human care. Miller et al. (2013) examined enucleated eyes from 70 pinnipeds, predominantly Zalophus californianus, but also from other species. Among the animals living in the wild, 35% (13 out of 37 individuals) exhibited an eye disease, while the prevalence was significantly higher, at 81.5%, in pinnipeds (22 out of 27 animals) in human care. Keratitis and pathological changes to the Descemet membrane were only documented in animals in human care. The study did not categorize individuals according to specific husbandry conditions (Miller 2013).

The aim of the present study is to investigate the relationship between the housing conditions of two groups of California sea lions — those housed in a freshwater facility and those in a saltwater facility at Nuremberg Zoo — and the prevalence of associated ocular diseases over a ten-year period. A primary advantage of comparing animals that reside in the same zoo is that they are exposed to similar environmental factors, such as sunlight duration and temperature as well as the same feeding regime and supplementation. The core objective of this study is to determine the extent to which water habitat may influence the incidence of ocular problems and other disease types in these animals, and to compare our findings with existing literature on the prevalence of eye diseases associated with different aquatic habitats.

## 2. Materials and Methods

### Data collection

We analyzed the medical records of 20 Californian sea lions aged at least 2 years old between 01.01.2012 and 31.12.2021. Initial descriptions and reports were made by the animal keepers. When an examination was possible, they were diagnosed and categorized by a veterinarian. Thus, our analysis differs from that of other authors who examined the individuals only once in different zoological institutions (Nakamura et al., 2021; Stach and Eule, 2021; Stach and Eule, 2024).

The diagnoses were divided into the categories “eye disease” (abbreviation eye), “injuries” (abbreviation inj), “skin disease” (abbreviation skin) and “other diseases” (abbreviation misc). The category “other diseases” includes rarely occurring diseases. The number of diagnoses in each category over the course of 10 years was counted (case or event). The category of eye diseases included “eye pinching” as a precursor of keratitis, corneal diseases (corneal opacity; keratitis), eye inflammation and cataract. If the diagnoses only indicated “cloudy eye” or “whitish eye”, this was assessed as corneal opacity. Lens opacity was only counted if the diagnosis was “cataract”.

The diagnoses were recorded in the data sheet as an event in the month when they were first registered in the medical record. If the same diagnosis occurred after more than 30 days without a clear connection to the previously recorded event, it was counted as a new diagnosis. This was also applied if, for example, one eye was initially affected by the same disease and after more than 30 days the second eye was affected, too. If two diagnoses relating to the same organ or organ system were mentioned in one month, the more serious finding was evaluated, e.g., if “squinting eyes” and “cloudy eyes” were mentioned, this was recorded as one event under “corneal opacity”. Duration of an illness or medical treatment could not in all cases be taken from the medical records, therefore this aspect was not addressed.

### Data analysis

As some of the individuals were moved between the facilities, died, or were newly added during the period under investigation, the number of animals in the facilities investigated was not constant over the entire period and in the monthly analysis. The number of animals kept at the same time varied between 6 and 7 individuals, in each of the two enclosures.

For this reason, the number of months that each individual spent under a particular housing condition was determined. If an individual was transferred from one enclosure to the other during a month, the month of life was attributed to the enclosure in which more than half of the days of that month were spent. To be able to compare the populations, the documented cases of diseases were evaluated as the percentage of the occurrence of the respective disease in relation to the total number of months that the individuals spent under this housing condition.

**Figure 2.**
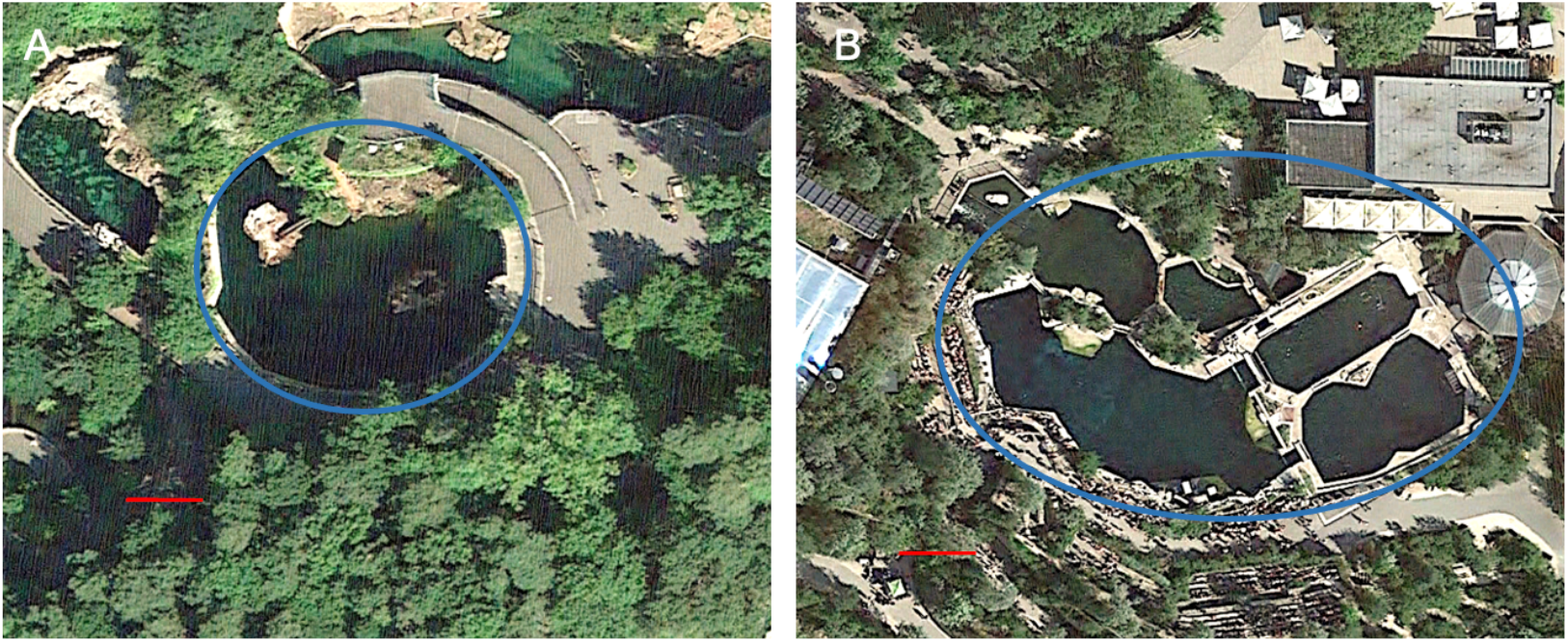
The two different facilities at Nuremberg Zoo where sea lions are kept, the blue circles indicate the pool areas. (**A**) Freshwater facility “Aquapark”. (**B**) Saltwater facility “Lagune”. The red line represents 10 m. Images: Google Earth, Landsat/Copernicus (23.08.2017).

### Housing conditions

Nuremberg Zoo has been keeping California sea lions since 1956. In March 2001 the enclosure “Aquapark” was opened. The outdoor enclosure consists of a 500 m2 land zone, and a pool with 840 m2 surface containing 2,3 million liters of freshwater. The pool has gray concrete walls which are non-reflective and a maximum depth of 3.5 m. Two land zones are integrated in the pool, of which one is the preferred hauling area of the sea lion group. A group of California sea lions and two female Harbor seals are housed under natural light and temperature conditions. Water temperatures vary between a minimum of 3°C in December until February and a maximum temperature of up to 24.5°C in July, depending on the air temperatures. Water treatment is done by filtering (drum and sand filter), phosphate precipitation, stabilization of the pH value and carbonate hardness by adding sodium hydrogen carbonate. There is also an around 100 m2 indoor area with cages to separate animals if necessary.

A second group of California sea lions was housed with bottlenose dolphins (*Tursiops truncatus*) in the saltwater facility “Lagune” (Lagoon), which was opened in 2011. The enclosure consisted of six outdoor pools of various sizes and depths (from 0.5 m to 7 m) an indoor area with a total water area of approximately 1900 m2, containing 7.5 million liters of saltwater. In contrast to the Aquapark the water temperature does not fall below 18°C in the winter months. Water treatment is done also done by filtering (drum and sand filter), phosphate precipitation, stabilization of the pH value and carbonate hardness by adding sodium hydrogen carbonate, water disinfection in the Lagoon is made by adding ozone and only to a very limited extent by chlorination. The saltwater was made of a mixture of salt brine and fresh water and had a salinity of 3.1%. Temperature, pH-value, carbonate hardness were measured in a daily protocol in both enclosures, in the Lagoon also salinity is measured daily and the microbiology of the water is monitored once a week.

## 3. Results

### 3.1. Population and population changes during the survey period

20 animals, which were at least two years old, lived from 01.01.2012 to 31.12.2021 in Nuremberg Zoo. Of these, 17 individuals were born in Nuremberg Zoo, three animals (B02013; B04567 and B01823) were born in other zoological institutions. The females B02013 and B04567 already came to Nuremberg in 1994 and 2002, while the male B01823 (2014) was transferred to Nuremberg Zoo during the study period at the age 20 months. The age of the sea lions examined ranged between 2 and 29 years. Of the 20 animals, 5 females died during the study period 2012 to 2021, with a mean age of 23 years 1 month (median 26 years 10 months; standard deviation 6 years 8 months). 26 pups were born during the study period, of which 5 females (3 in 2013; 2 in 2017) remained in Nuremberg Zoo, all other pups were transferred to other zoos. One male (B01823) was added to the group in the Lagoon in 2014. All diagnoses of these 6 individuals (age ⩾2 years) are included in the analysis. During the survey period, 145 diagnoses were registered, with 29% (42 diagnoses) relating to eye diseases. 32% of the diagnoses concerned injuries, which were often bites, and the category of other diagnoses counts for another 32% of all cases. Skin diseases were diagnosed in 7% (10 cases). Individual B05798 (*12.06.1994; 23.03.2021) showed 38 cases of eye diseases, 1 case of skin disease, 5 injuries, and 13 other diagnoses during her lifetime, which adds up to 57 diagnoses. Thus, she had more diseases than any other individual in this study. We excluded this individual which was housed at the salt water facility from the data we show in the following.

The exclusion of B05798 from the data analysis reduces the number of diagnoses in the period 2012 to 2021 to 129 diagnoses in 19 animals. Of these, 25 % (i.e. -4 %; 32 diagnoses) are attributable to eye diseases, 8 % (10 diagnoses) to skin diseases, 33 % (42 diagnoses) to injuries and 35 % (45 cases) to miscellaneous diagnoses. In Figure 3 we plotted the occurrence of the different diagnoses corrected for the time animals spend in the saltwater or freshwater facility respectively. Injuries were the most common diagnosis (42 cases), followed by eye diseases (32 cases) and skin diseases (10 cases). The category of other diagnoses included gastrointestinal diagnoses, reduced food intake, but also behavioral abnormalities (10 cases).

**Figure 3.**
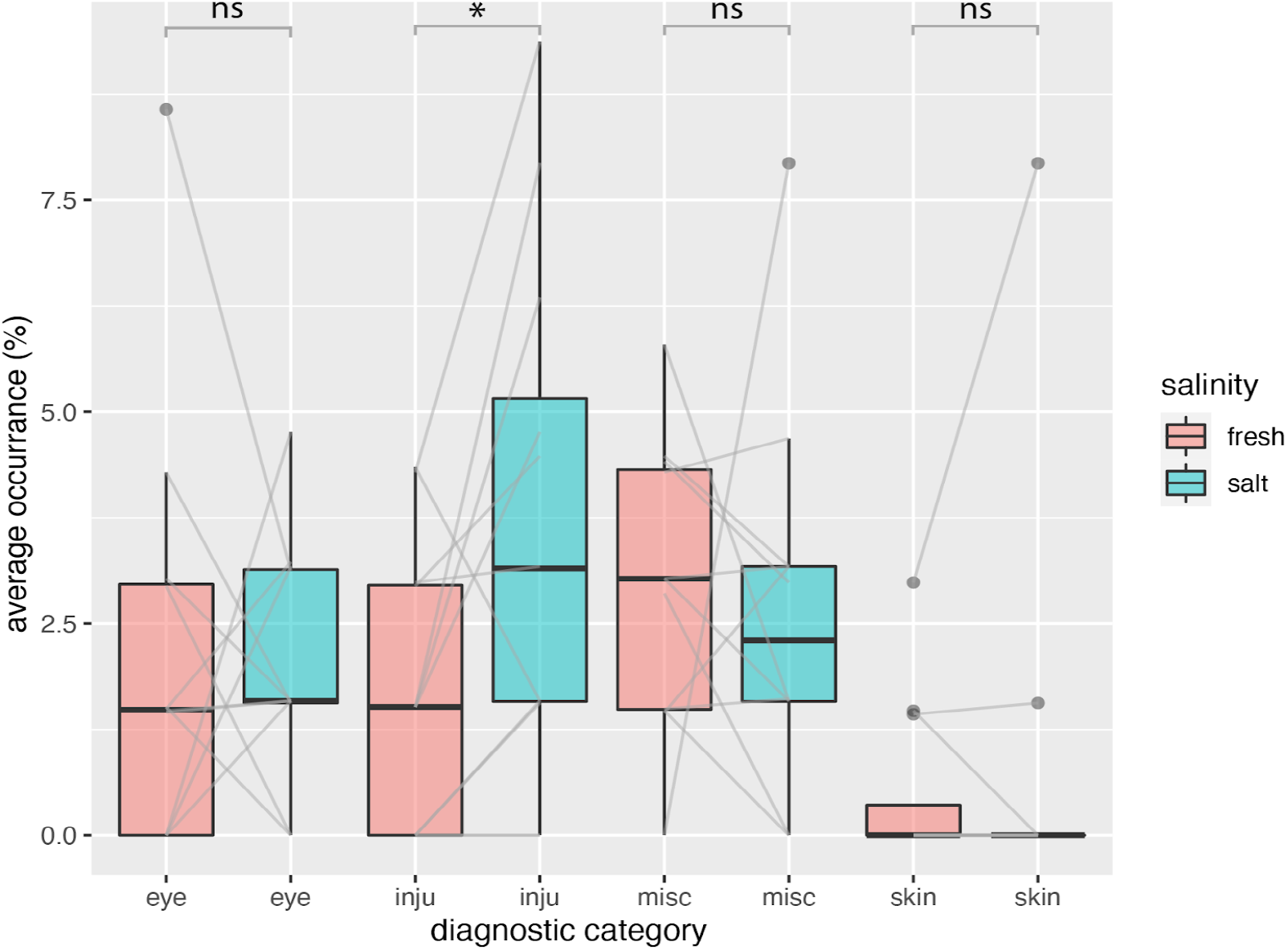
Average occurrence of certain diagnostic findings for the fresh- (red) and saltwater (green) facilities. For each month, we calculated the average occurrence of specific diagnoses over a ten-year period, adjusted for the total number of months each individual lived in the facility. Each boxplot displays the variation of the occurrences for the different month of a year which are connected with a gray line in between the fresh- and saltwater category. Note that some lines are double, which is not always visible. X-axis labels: eye = eye disease, inju = injuries, skin = skin disease and misc = other diagnoses.

### 3.2. Occurrence of diagnoses

We calculated the occurrence of specific diagnoses (expressed as a percentage per month over a ten-year period) based on the time the animals spent in each facility. For each month of the year, we derived average occurrences of various diagnostic categories, averaged over ten years (Figure 3). We compared these occurrences across all months and diagnostic categories and found no significant difference between the saltwater and freshwater facilities for eye-related diseases (Wilcoxon Signed-Rank test; freshwater (Mdn = 1.5, n = 12), saltwater (Mdn = 1.6, n = 12), W+ = 42, p = 0.850, r = 0.05). Similarly, there was no significant difference in miscellaneous diagnoses between the freshwater (Mdn = 3, n = 12) and saltwater (Mdn = 2.3, n = 12) facilities (W+ = 26, p = 0.339, r = -0.3). However, we did find a significant difference in injuries between the freshwater (Mdn = 1.5, n = 12) and saltwater (Mdn = 3.1, n = 12) facilities (W+ = 49, p = 0.027, r = 0.7). The skin disease category had too few samples for a Wilcoxon Signed-Rank test.

To further explore whether the occurrence of different diseases varied by month, we conducted a chi-square test of independence. Figure 4A indicates that skin diseases are most frequently diagnosed in January, eye diseases peak in July, and injuries occur often in February. However, the residuals from expected values were comparatively low (maximum = 2.45), therefore there was no significant association between diagnostic category and month for the freshwater facility (X^2^ = 33.097, df = 33, p = 0.4625). Figure 4B reveals that skin diseases are predominantly diagnosed in January (similar to the freshwater facility), while eye diseases show small peaks in November and May. Because the residuals in this case were comparatively high (maximum = 4.87), we found a significant association between diagnostic categories and month for the saltwater facility (X^2^ = 67.261, df = 33, p = 0.0003934). When cumulative occurrences of diagnostic categories were directly compared between the freshwater and saltwater facility (see Figure 4C), it became evident that injuries are overrepresented in the saltwater facility, while eye diseases are somewhat above the expected value in the freshwater facility. Despite this observation, the residuals were also relatively small (maximum = 1.38), resulting in no significant association (X^2^ = 7.6069, df = 3, p = 0.05487).

**Figure 4.**
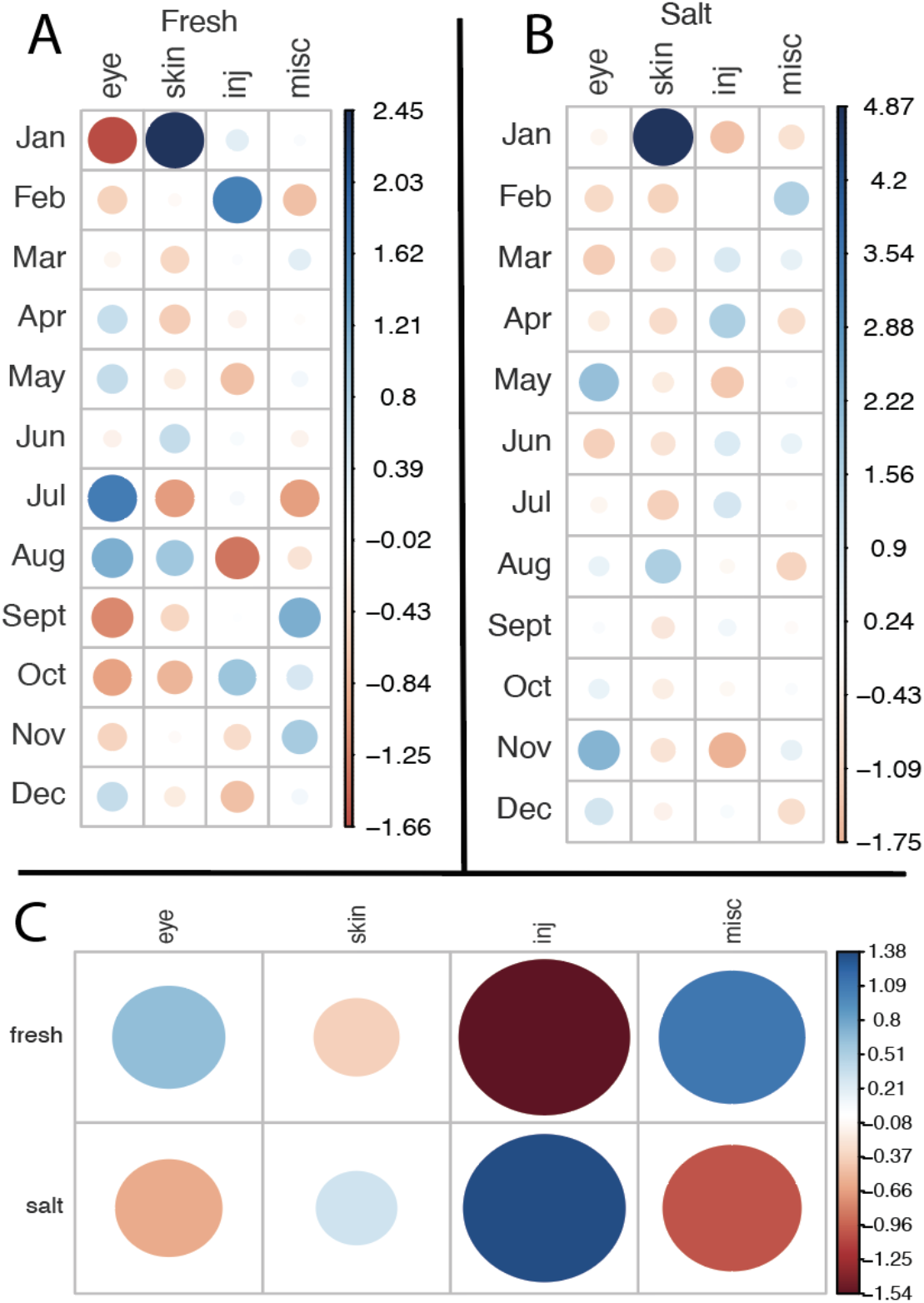
Pearson residuals from the expected values for the average occurrences (percent per month) of certain diagnostic categories. Values are based on the time the animals lived in the respective facility. Positive residuals are indicated in blue and specify an attraction (positive association) between the corresponding row and column variables. Negative residuals are indicated in red and imply a repulsion (negative association) between the corresponding row and column variables. The size of the circle is proportional to the amount of the cell contribution. (A) Residual plot for the fresh water facility. (B) Residual plot for the salt water facility, note that the residuals are very high compared to the other plots. (C) Residual plot directly comparing salt and fresh water facilities.

**Figure 5.**
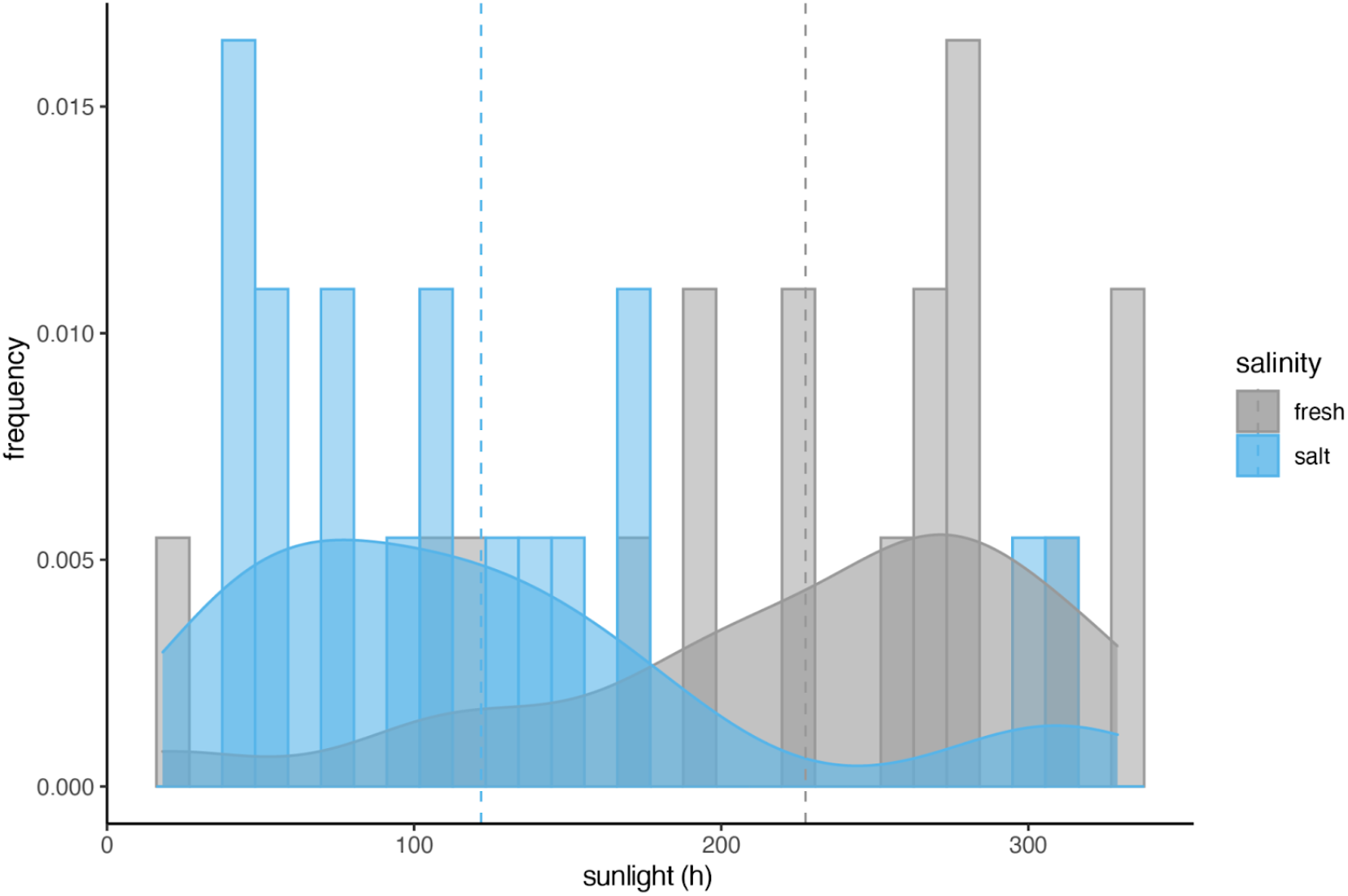
Frequency of ocular diseases during different amounts of hours of sunshine. The saltwater facility is indicated in blue; the freshwater facility is indicated in gray. The dashed lines indicate the mean values. If UV radiation had an influence on the occurrence of ocular diseases, there should be a similar distribution for fresh and saltwater.

### 3.3. Occurrence of ocular diseases

We also tested if the ocular diseases were occurring more frequently during times where sunlight hours were higher. Interestingly we found a significant difference in the frequency of ocular diseases between fresh and saltwater (Wilcoxon Signed-Rank test; W+ = 16, p = .003, r = -0.7). Ocular diseases were diagnosed more often during periods with higher sunlight hours for the freshwater facility (Mdn = 256.5), whereas ocular diseases were diagnosed more often during periods with less light for the saltwater facility (Mdn = 106.2).

4. Discussion

In this study, we investigated the prevalence of ocular diseases in California sea lions at Nuremberg Zoo, focusing on the influence of water habitat - freshwater versus saltwater. During our 10 year survey period between 01.01.2012 and 31.12.2021 we recorded 32 cases of eye diseases. These accounted for 25 % of 129 cases of medical records in total for 19 individuals investigated. Our results indicate that salinity does not significantly affect the incidence of ocular diseases in sea lions. Although we observed a trend towards an increase in eye problems during the summer months in the freshwater habitat, this trend was not statistically significant. The increase is likely to be related to deteriorating water quality due to higher temperatures and longer daylight hours. In contrast, the saltwater facility features a superior filtration and life support system, which ensures more stable water quality throughout the year. However, the total amount of ocular diseases did not significantly differ between the salt and freshwater facility. The analysis of sunlight exposure and its correlation with the frequency of ocular diseases revealed distinct patterns between the freshwater and saltwater facilities. As all sea lions were housed at Nuremberg Zoo, they experienced the same climate, were subjected to the same amount of sunlight and also received the same diet and supplementation. Therefore, one might expect a similar trend in the frequency of ocular diseases in both freshwater and saltwater facilities if UV radiation had a significant influence. However, it is important to note that the freshwater facility is considerably smaller, with less opportunities to rest on land, which may result in less shaded areas for the animals. We did not assess the specific amount of shade or the locations where the sea lions rested or were fed throughout the day, which could provide insights into their exposure to sunlight. This lack of data may limit our understanding of the full impact of UV radiation on ocular health in these animals. Future studies should consider evaluating the shading patterns and resting behaviors of the sea lions to better understand the relationship between sunlight exposure and ocular disease prevalence.

Eye problems in sea lions are multifactorial, as suggested by Colitz et al. (2022; 2019) and other authors (Nakamura et al., 2021; Gage, 2011), therefore we believe it would be a significant oversight for guidelines to focus exclusively on the freshwater versus saltwater debate. Although the current study is preliminary and not yet conclusive, it suggests that ocular diseases occur with similar frequencies in both saltwater and freshwater facilities. We strongly suggest that further research is needed to address this complex issue. A meaningful comparison of ocular conditions in sea lions living in saltwater and those living in freshwater environments should be made, taking into account a wide range of variables beyond salinity.

Key parameters to consider include water quality, disinfection methods, pH levels, temperature and pool paint materials, as well as environmental factors such as seasonal changes, sun exposure and UV radiation, also same diet and supplementation. Regular monitoring of these factors over a 12-month period is essential to establish correlations between environmental conditions and eye health. This should be combined with detailed ocular monitoring by veterinarians, including photographic documentation, to track the condition of the eyes over time rather than just assessing them when a disease is diagnosed, as it was made for this study.

We propose a systematic approach to photographing the eyes of sea lions on a monthly basis. This process will involve the implementation of a fixed head position, alongside a defined camera angle and distance, to ensure consistency in each image captured. To mitigate issues such as light reflections on the eye, we emphasize the importance of controlling lighting conditions. Conducting the photography sessions in indoor enclosures will allow for the use of indirect lighting, which helps to diffuse harsh light and reduces glare. By carefully selecting the type of lighting and positioning it in a way that uniformly illuminates the head of the sea lion, it will be possible to standardize the procedure. This will enhance image clarity by minimizing unwanted reflections and shadows, leading to more reliable diagnostic results. In addition to the fixed head positioning and controlled lighting, we recommend developing a standardized protocol to accommodate the animals prior to imaging. This includes familiarizing them with the setup as part of medical training sessions, which will ensure the animals are comfortable during the process, making regular imaging sessions more efficient and effective. By adhering to these methods, we aim to create a robust standard operating procedure that minimizes variability in image capture, thus facilitating accurate assessment and diagnosis of any ocular conditions present in the sea lions, while contributing to the overall welfare of the animals involved in the study.

We propose a holistic approach to managing the eye health of pinnipeds. Preventive strategies should include improving water filtration, closely monitoring microbial influences, measure light exposure and provide shaded areas. Without a comprehensive approach, ocular diseases are likely to persist in managed populations. In conclusion, it is essential to shift the focus from a rather opinion based freshwater versus saltwater debate to a comprehensive investigation based on factual evidence. This should include an examination of all potential contributing factors, such as consistent veterinary eye examinations and photographic documentation, to ensure an accurate assessment of disease progression.

## Author Contributions

Conceptualization, I.B., S.H., K.B. and L.v.F.; methodology, I.B., S.H., K.B. and L.v.F.; validation, I.B., S.H. and R.S.; formal analysis, I.B., S.H. and R.S.; investigation, I.B., S.H. and R.S.; resources, K.B. and J.B.; data curation, I.B., S.H. and R.S.; writing—original draft preparation, I.B., R.S., J.B. and L.v.F.; writing—review and editing, I.B., R.S., J.B. and L.v.F.; visualization, R.S.; supervision, I.B., L.v.F. and K.B.; project administration, L.v.F., J.B. and K.B.; All authors have read and agreed to the published version of the manuscript.

## Institutional Review Board Statement

Ethical review and approval were waived for this study because observations only took place during regular or mandatory veterinary observations.

## Acknowledgments

We would like to thank all the animal caretakers from the Lagoon and Aquapark facilities, especially their leads, Armin Fritz and Thorsten Krist, for their dedication, time, and effort in closely monitoring the health of the sea lions. Their commitment to animal welfare has been invaluable to our research. We also extend our appreciation to the veterinary staff for their continuous support and expertise throughout the study.

## References

BMEL (Bundesministerium für Ernährung und Landwirtschaft/German Federal Ministry of Food and Agriculture) 2020, Gutachten über die Mindestanforderungen an die Haltung von Säugetieren, p. 192 ff, Berlin.

De Haan K (2011) Corneal lesions in captive California Sea Lions (Zalophus californius) The diagnostic aid of fluorescein and the influence of water quality on the cornea. Utrecht University. https://studenttheses.uu.nl/handle/20.500.12932/9560

Dunn JL, Overstrom NA, St Alibis DJ, Abt DA (1996) An epidemiologic survey to determine factors associated with corneal and lenticular lesions in captive harbor seals and California sea lions. IAAAM, Poster abstract.

Colitz CM, Saville WJ, Renner MS, McBain JF, Reidarson TH, Schmitt TL, Nolan EC, Dugan SJ, Knightly F, Rodriguez MM, Mejia-Fava JC, Osborn SD, Clough PL, Collins SP, Osborn BA, Terrell K. (2010a) Risk factors associated with cataracts and lens luxations in captive pinnipeds in the United States and the Bahamas. J Am Vet Med Assoc.; 237(4): 429–36. doi: 10.2460/javma.237.4.429.

Colitz CM, Renner MS, Manire CA, Doescher B, Schmitt TL, Osborn SD, Croft L, Olds J, Gehring E, Mergl J, Tuttle AD, Sutherland-Smith M, Rudnick JC. (2010b) Characterization of progressive keratitis in otariids. Vet Ophthalmol.; 13 Suppl: 47–53. doi: 10.1111/j.1463-5224.2010.00811.x.

Colitz CM, Saville WJA., Walsh MT, Latson E (2019). Factors associated with keratopathy in captive pinnipeds. Journal of the American Veterinary Medical Association, 255(2), 224–230. 10.2460/javma.255.2.224

Colitz CM (2022). Ophthalmology of Pinnipedimorpha: Seals, Sea Lions, and Walruses. In: Montiani-Ferreira, F., Moore, B.A., Ben-Shlomo, G. (eds) Wild and Exotic Animal Ophthalmology. Springer, Cham. 10.1007/978-3-030-81273-7_13

Enns, A., Kraus, D., & Hebb, A. (2020). Ours to save: the distribution, status and conservation needs of Canada’s endemic species. NatureServe Canada and Nature Conservancy of Canada, 75.

Gage LJ (2009). Overview of Known and Suspected Causes of Captive Pinniped Eye Problems. IAAAM, Poster abstract.

Gage LJ (2011) Captive pinniped eye problems, we can do better! Journal of Marine Animals and Their Ecology, Vol. 4, No 2, 2011, 25–28.

Gulland FMD, Dierauf LA, Whitman KL (Eds.). (2018). CRC Handbook of Marine Mammal Medicine (3rd ed.). CRC Press. 10.1201/9781315144931

Harrison J (2018). Sea Lions in the Willamette. Northwest Power and Conservation Council. Retrieved July 24, 2023. https://www.nwcouncil.org/news/sea-lions-willamette/

Hauser D. D., Allen, C. S., Rich Jr, H. B., & Quinn, T. P. (2008). Resident harbor seals (Phoca vitulina) in Iliamna Lake, Alaska: summer diet and partial consumption of adult sockeye salmon (Oncorhynchus nerka). Aquatic Mammals, 34(3), 303.|DOI 10.1578/AM.34.3.2008.303

Latson E. (2011) By-Products of Disinfection of Water and Potential Mechanisms of Ocular and Other Organ Injury and Hemosiderosis in Marine Mammals, IAAM, Poster abstract.

Miller S, Colitz CM, St Leger J, Dubielzig R. (2013) A retrospective survey of the ocular histopathology of the pinniped eye with emphasis on corneal disease. Vet Ophthalmol.; 16(2):119–29. doi: 10.1111/j.1463-5224.2012.01040.x.

Nakamura M, Matsushiro M, Tsunokawa M, Maehara S, Kooriyama T (2021) Survey of ophthalmic disorders among captive pinnipeds in Japan. J Vet Med Sci.; 83 (7):1075–1080. doi: 10.1292/jvms.20-0329.

Parsons EC (2013). An Introduction to Marine Mammal Biology and Conservation. Jones & Bartlett. p. 118. ISBN 978-0-7637-8344-0.

Smith, RJ; Cox, TM; Westgate, AJ (2006). Movements of Harbor Seals (Phoca Vitulina Mellonae) in Lacs des Loups Marins, Quebec. Marine Mammal Science. 22 (2): 485.

Stach MR, Eule CJ (2021) Epidemiological Survey on Ocular Diseases of Pinnipeds—Period Prevalence and Husbandry Conditions within Central European Facilities, IAAAM, Poster abstract.

Stach MR, Eule CJ (2024) Pinniped species in German facilities: Point prevalence of ocular disorders, Veterinary Ophthalmology, Poster abstract, 10.1111/vop.13253

Stewart JE, Pomeroy PP, Duck CD, Twiss SD, (2014). Finescale ecological niche modeling provides evidence that lactating gray seals (Halichoerus grypus) prefer access to fresh water in order to drink. Mar Mam Sci, 30: 1456–1472. 10.1111/mms.12126

Picture in Figure 1 by Sergey Gabdurakhmanov from Mountain View, USA - Nerpa (Pusa sibirica), CC BY 2.0, https://commons.wikmedia.org/w/index.php?curid=132821056

